# There and back again – unraveling mechanisms of bacterial biogeography in the North Pacific Subtropical Gyre to and from station ALOHA

**DOI:** 10.1101/141085

**Authors:** Markus V. Lindh

## Abstract

Bacterially-mediated fluxes of energy and matter are dynamic in time and space coupled with shifts in bacterial community structure. Yet, our understanding of mechanisms shaping bacterial biogeography remains limited. Near-surface seawater was collected during transits between Honolulu and Station ALOHA in the North Pacific Subtropical Gyre to examine the shape of occupancy-frequency distributions (the different number of populations occupying different number of sites) and determine bacterial metapopulation dynamics. Bacterial 16S rRNA gene amplicons were sequenced from whole seawater and filter-size fractionated plankton DNA samples while also separating the community into distinct taxonomic groups at phyla/class and analyzing these compartments separately. For the total seawater (i.e. the >0.2 μm size fraction) and picoplankton communities (i.e. the size fraction >0.2 μm and < 3.0 μm), but not the large size fraction community (i.e. the >3.0 μm size fraction), most individual operational taxonomic units (OTUs) occupied a single site and the number of OTUs occupying different number of sites followed a significant bimodal pattern with several core OTUs occupying all sites. Nevertheless, only Cyanobacteria (in particular *Prochlorococcus* sp.) and in a few instances also Alphaproteobacteria (in particular SAR11 clade and Aegan-169 marine group bacteria) exhibited bimodal occupancy-frequency patterns. As expected, *Prochlorococcus* sp. had an inversed bimodal occupancy-frequency distribution with most OTUs found at all sites. Yet, there were individual satellite OTUs affiliated with *Prochlorococcus* sp. that were phylogenetically distinct from the core OTUs and only found at a single site. Collectively, these findings indicate that different compartments (size fractions and taxa) have different metapopulation dynamics. Bimodal patterns among the low diversity total and picoplankton communities but not in the high diversity large size fraction suggest that positive feedbacks between local abundance and occupancy are important when environmental conditions are homogenous and diversity is low.

## Introduction

Spatial and temporal variability in bacterial population dynamics influence nutrient cycling in aquatic systems (Crump et al., 2004; Kirchman et al., 2005; Fuhrman et al., 2006; Galand et al., 2010; Lindström et al., 2010; Östman et al., 2010; Alonso-Saez et al., 2015). Yet, despite observable biogeographical patterns among marine microbial assemblages (see e.g. Pommier et al., 2007; Ghiglione et al., 2012; Sunagawa et al., 2015; Salazar et al., 2016), the mechanisms shaping microbial biogeography remain largely unknown (Martiny et al., 2006; Hanson et al., 2012; Poisot et al., 2013).

With the introduction of high-throughput sequencing, and large sequence datasets enabled by these technologies, microbial ecologists are now able to test a wide variety of theoretical ecological models that are the foundation for mechanisms explaining macroecological patterns among larger taxa (Purdy et al., 2010; Poisot et al., 2013). Such theoretical models, including, but not limited to, metacommunity (Leibold et al., 2004) and metapopulation (McGeoch and Gaston, 2002) frameworks have provided mechanistic concepts in ecology among organisms ranging from birds to fish and insects to phytoplankton (Levin, 1974; Hanski, 1982; Gotelli, 1991; Tokeshi, 1992; Hanski and Gyllenberg, 1993; van Rensburg et al., 2000; Hubbell, 2001; Mehranvar and Jackson, 2001; Mouquet and Loreau M, 2002; Verberk et al., 2010; Unterseher et al., 2011; Wardle et al., 2011; Hercos et al., 2013).

Two models; Hanski’s core and satellite hypothesis (CSH; (Hanski, 1982) and Levin’s model (Levin, 1974), were recently empirically tested and used to explain distribution patterns among marine bacteria that are typically assumed not to be dispersal limited (Lindh et al., 2016). In the study bimodal occupancy-frequency patterns (i.e. the number of species occupying different number of sites) were found and the CSH model agreed well with observed data in the Baltic Sea and in global datasets, indicating its applicability also for marine microbes. The CSH model provides a mechanistic basis for the observation of abundant (core) and rare (satellite) species in ecosystems. In brief, the CSH makes predictions of the shape of occupancy-frequency distributions from variation in colonization and extinction rates (i.e. populations successfully dispersed to previously unoccupied sites vs. populations disappearing from previously occupied sites). In the CSH the bimodal occupancy-frequency distributions are the result of stochastic variation in colonization and extinction rates (i.e. a quadratic relationship between colonization/extinction rates and occupancy) that either push populations to become rare or abundant. Nevertheless, empirical data from examining the applicability of metapopulation models to marine bacterial assemblages are limited.

In the present paper the prevalence of bimodal occupancy-frequency patterns were examined in the oligotrophic North Pacific Ocean and among different taxa and/or size fractions of a bacterial community. The main hypothesis was that in this relatively homogenous oligotrophic ocean environment bimodal patterns, reflecting a coherent oceanic region without dispersal limitation and environmental filtering, could be a common feature among marine bacteria. The second hypothesis was that taxa with different dispersal capability and filtered by various environmental conditions could display different metapopulation dynamics. The third hypothesis was that since *Prochlorococcus* sp. is the dominant bacteria found in this system (Schmidt et al., 1991; Campbell and Vaulot, 1993; Eiler et al., 2011) much of the observed metapopulation dynamics could be driven by this key organism but that there may be core- and satellite distribution patterns within the same genus. To test these hypotheses, bacterial 16S rRNA amplicons were sequenced from whole seawater and filter size fractionated community DNA, sampled from the near-surface ocean on a series of cruise transects between Honolulu and station ALOHA (22.75° N, 158° W) in the North Pacific Subtropical Gyre. The observed bacterial populations were subsequently fitted to different theoretical metapopulation models.

## Material and Methods

### Field sampling

Seawater samples for subsequent extraction of plankton DNA were collected during 4 research cruises (October 2015, HOT 277; November 2015, KM1519; December 2015, KM1521; and January 2016, HOT 280) offshore of the Hawaiian island of Oahu aboard the R/V Ka’imikai-O-kanaloa and R/V *Kilo Moana.* Seawater was collected from the research vessels’ flow-through seawater intake systems into acid-washed, Milli-Q rinsed polycarbonate bottles. The flow-through seawater system was instrumented to include a thermosalinometer (SeaBird 911), fluorometer (Seapoint Chlorophyll Fluorometer; Seapoint Sensors, Inc.) and dissolved oxygen (O_2_) sensor (SBE 43; Sea-Bird Electronics). On all cruises, plankton biomass for subsequent DNA extraction was harvested by filtering 2 L of seawater on to 25 mm diameter, 0.2 μm pore size polyethersulfone filters (Supor membrane, Pall). In addition, on two of the cruises (December 2015 and January 2016) plankton biomass were also sequentially filter size-fractionated samples for subsequent extraction of DNA for information on bacterial community structure among different plankton size classes. During these cruises, seawater was filtered onto 25 mm diameter, 3.0 μm pore size polycarbonate membranes (Millipore), followed by filtration onto 25 mm diameter, 0.2 μm pore size Supor filters. The various filter size-fractionated bacterial communities were defined as: total (>0.2 μm), picoplankton (>0.2 and <3 μm) and large (>3 μm), respectively. Filters were stored at - 80°C until extraction.

Coincident samples for measurement of bacterial abundance and production were collected from the vessels’ flow-through seawater system. Seawater for determination of bacterial abundance was subsampled into 2 ml cryotubes (BD Falcon) preserved with 20 μL paraformaldehyde (Final concentration 0.2%; Sigma-Aldrich). Triplicate 1.5 ml samples for subsequent measurements of ^3^H-leucine incorporation rates were aliquoted into 2.0 ml microcentrifuge tubes (Axygen; (Pace et al., 2004)) and amended with 20 nM (final concentration) of ^3^H-leucine stock (3, 4, 5 -^3^H-leucine, 109 Ci/mmol; Perkin-Elmer). One killed control was included from each station sampled (5% final concentration of trichloroacetic acid; Sigma-Aldrich). Samples for ^3^H-leucine incorporation rates were incubated shipboard at *in situ* temperatures following the ^3^H-leucine incorporation protocol described in (Kirchman et al., 1985) and (Smith and Azam, 1992). Preserved bacterial abundance and killed ^3^H-leucine incorporation rate samples were flash frozen in liquid nitrogen and stored at -80° C until processing in the laboratory. Preserved bacterial abundance samples were thawed, and aliquoted into 250 μl wells and stained with 2.5μl μl of 100X SybrGreen (Thermo Fisher) and enumerated using an Attune™ acoustic focusing flow cytometer (Applied Biosystems). ^3^H-leucine incorporation rates samples were processed following a modified method of the microcentrifuge method as described in (Smith and Azam, 1992). A detailed description of the ^3^H-leucine incorporation protocol can be found in (Viviani and Church, 2017).

### DNA extraction, PCR, sequence processing and analysis

DNA was extracted from the filters using the DNeasy Plant MiniKit (Qiagen) following slight modifications of the manufacturer’s suggestions. Modifications included the addition of Proteinase K to the lysis buffer followed by bead-beating with 0.1 mm and 0.5 mm glass beads (Biospec products) for additional cell disruption prior to the extraction. DNA concentrations were determined using the Qubit 2.0 Fluorometer and Qubit dsDNA High Sensitivity Assay kit (Molecular Probes), with DNA quality checked using 1.5% agarose gel electrophoresis. Extracted DNA was stored at −80°C.

Amplicon processing for all samples was performed as described in (Lindh et al., 2015). Bacterial 16S rRNA was amplified using primers 341F and 805R (Herlemann et al., 2011) following the PCR protocol of (Hugerth et al., 2014). Amplicons were purified by spin-column centrifugation using E.Z.N.A.^®^ Cycle Pure kit (Omega Biotek). The resulting purified amplicons were pooled in equimolar concentration and sequenced on an Illumina Miseq (Illumina, USA) platform at the Hawai’i Institute for Marine Biology (HIMB), Hawaii, USA using the 300 bp paired-end setting. Raw sequence data generated from Illumina Miseq were processed using the UPARSE pipeline (Edgar, 2013). Taxonomy was determined against the SINA/SILVA database (SILVA123; (Quast et al., 2013)). After quality filtering and discarding plastid and archaeal sequences a total of 1.25 million bacterial sequences were utilized for subsequent analyses. Thus, the final OTU table consisted of 96 samples with 27340 OTUs delineated at 99% 16S rRNA gene identity with an average of 13093±5644 sequences per sample. For all alpha diversity measures the OTU table were subsampled to 10,000 sequences per sample. DNA sequences have been deposited in the National Center for Biotechnology Information (NCBI) Sequence Read Archive under accession number SRP091841.

### Statistical tests

Occupancy-frequency distributions (the number of OTUs occupying different number of sites) were analyzed as described in (Lindh et al., 2016). In brief, an equivalent to Tokeshi’s test of bimodality was performed using Mitchell-Olds’ and Shaw’s test (Mitchell-Olds and Shaw, 1987) for the location of quadratic extremes. Colonization and extinction rates of OTUs were determined by calculating the change in fraction of sites occupied between two cruises where the same stations were sampled with < 1 month interval. Non-linear least squares analysis of observed colonization and extinction rates was performed using the equations of Levin’s model, the CSH hypothesis and Gotelli’s propagule rain model (Levin, 1974; Hanski, 1982; Gotelli, 1991).

All statistical tests were performed in R 3.3.3 (Team, 2014) using the package “Vegan” (Jari Oksanen et al., 2010). Graphical outputs were made in R 3.3.3 using the package “ggplot2” (Wickham, 2009).

## Results

In the present study bacterial population dynamics and biogeography were examined during 4 research cruises in the North Pacific Subtropical Gyre (Fig. 1). For all four transects (total 96 samples) 27,340 operational taxonomic units (OTUs) were detected. A more exhaustive sampling was performed in the October 2015 cruise with measurements of a larger set of biotic and abiotic measurements and is thus the focus in the present paper. An additional cruise with a different transect (November 2015) to the Southwest of Oahu were performed to compare patterns observed in the October 2015 cruise. Finally, on two cruises (December 2015 and January 2016), samples were filter size-fractionated for information on bacterial diversity associated with differing size classes of plankton (>3 μm, >0.2 and <3.0 μm, and >0.2 μm, large, picoplankton and total, respectively).

**Figure 1.**
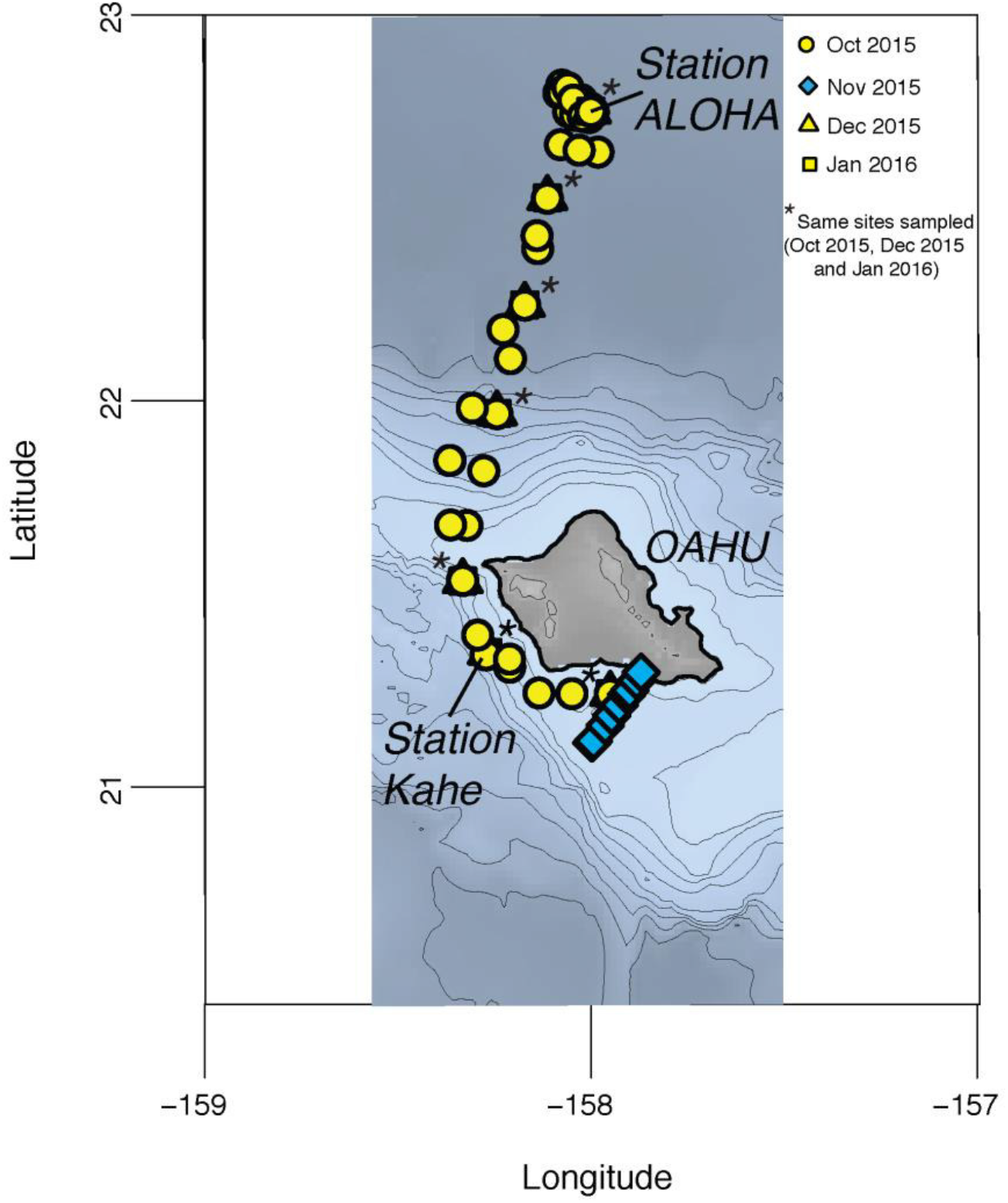
Map illustrating the four transects performed with the locations of Station ALOHA and Station Kahe marked. Asterisks denote the same station sample repeatedly.

## Physicochemical conditions, community abundance, production and biodiversity

Across the 200 km transect to Station ALOHA (October 2015) concentrations of Chl *a* and sea surface temperatures were relatively stable in space, averaging 0.6 μg L^−1^ and 26°C, respectively (Fig. 2). Bacterial abundance varied ~2-fold (ranging 5.73 x10^5^ cells ml^−1^ to 9.13 x 10^5^ cells ml^−1^; Fig. 2). ^3^H-leucine incorporation rates ranged between 13 and 37 pmol Leu L^−1^ h^−1^ (Fig. 2). Variation in bacterial abundance and production was overall small (SD = 0.7 x10^5^ cells ml^−1^ and SD = 6 pmol Leu L^−1^ h^−1^; Fig. 2).

**Figure 2.**
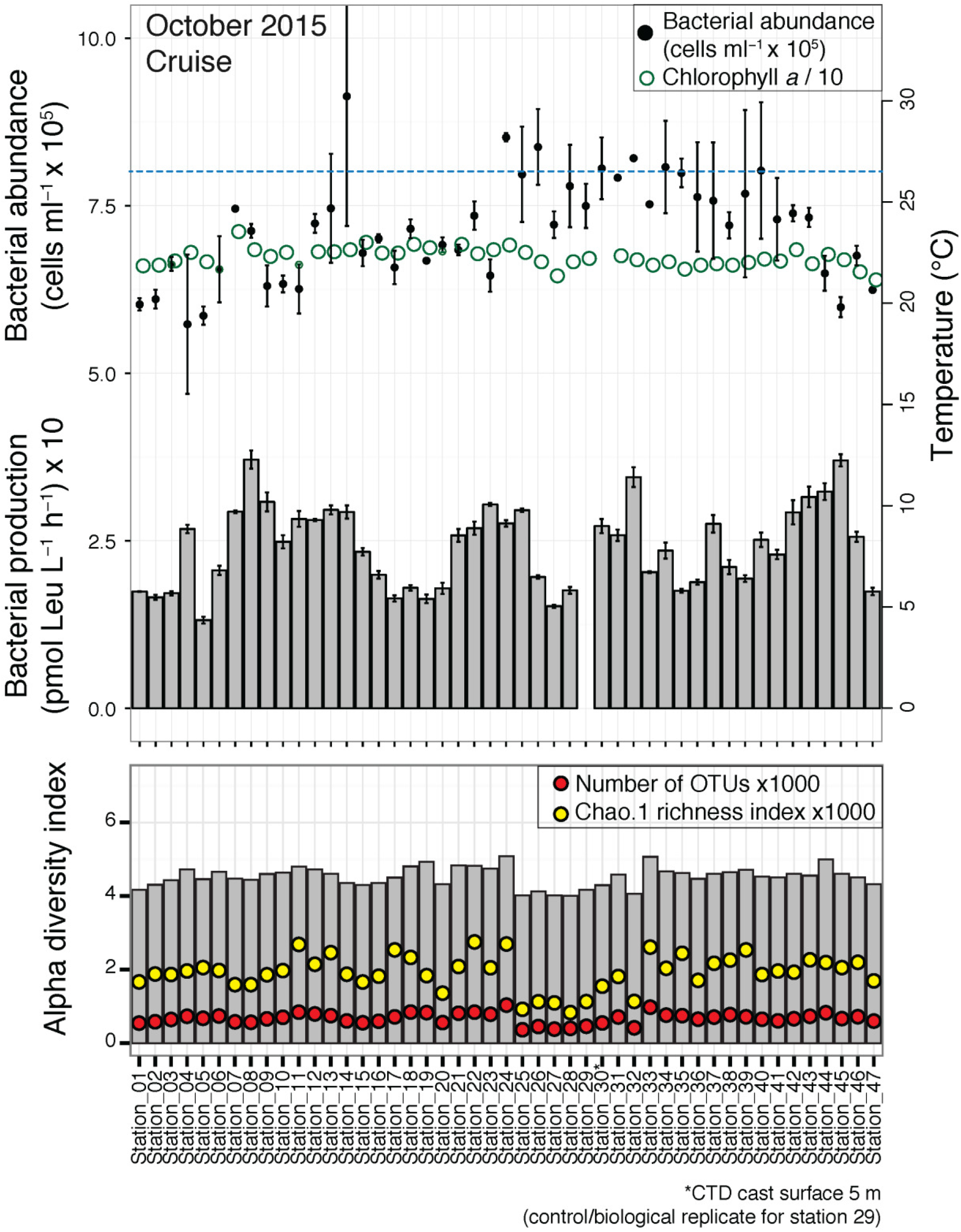
Variation bacterial community abundance, production and alpha diversity during the October 2015 cruise, including variation in Chl *a* (green open circles) and average sea surface temperature (blue dotted line).

The number of OTUs varied between 371 and 1037 but appeared relatively stable in space and time (SD = 145.34). Estimation of alpha diversity followed a similar trend where Shannon diversity index and Chao 1 richness varied between 3.99 and 5.06and 903.45 to 2256.88, respectively. Alpha diversity for the total community was also stable within all other cruises around the same range as observed during October 2015 (Fig. S1A). During the December 2015 and January 2016 transect, size fractionation of plankton DNA samples revealed higher alpha diversity in the larger size fraction (3.0 μm filter fraction) compared to the total (>0.2 μm) and picoplankton fraction (0.2 and 3.0 μm; Fig. S1A).

## Bacterial community composition

In all cruises the total and picoplankton communities were dominated by Cyanobacteria (averaging ~50% of total sequences), with bacteria that were affiliated with several other phyla or classes not included in the taxonomic analysis or unclassified OTUs comprising nearly 25% of the total sequences (Fig. 3). Actinobacteria and Alphaproteobacteria each typically contributed to ~5% of total sequences in these communities. In the December 2015 and January 2016 cruises the large size class bacterial assemblages were more diverse than other size classes of bacteria, with notable increases in relative abundances of Alphaproteobacteria, Planctomycetes, Chloroflexi and Bacteroidetes (Fig. 3B). However, Cyanobacteria were also dominating the large size class communities. Cluster analysis of Bray-Curtis distances obtained from samples in the October 2015 cruise showed a largely similar community structure (~15% dissimilarity), resulting in samples from near-shore coastal sites clustering with samples from the open ocean sites (Fig. 3).

**Figure 3.**
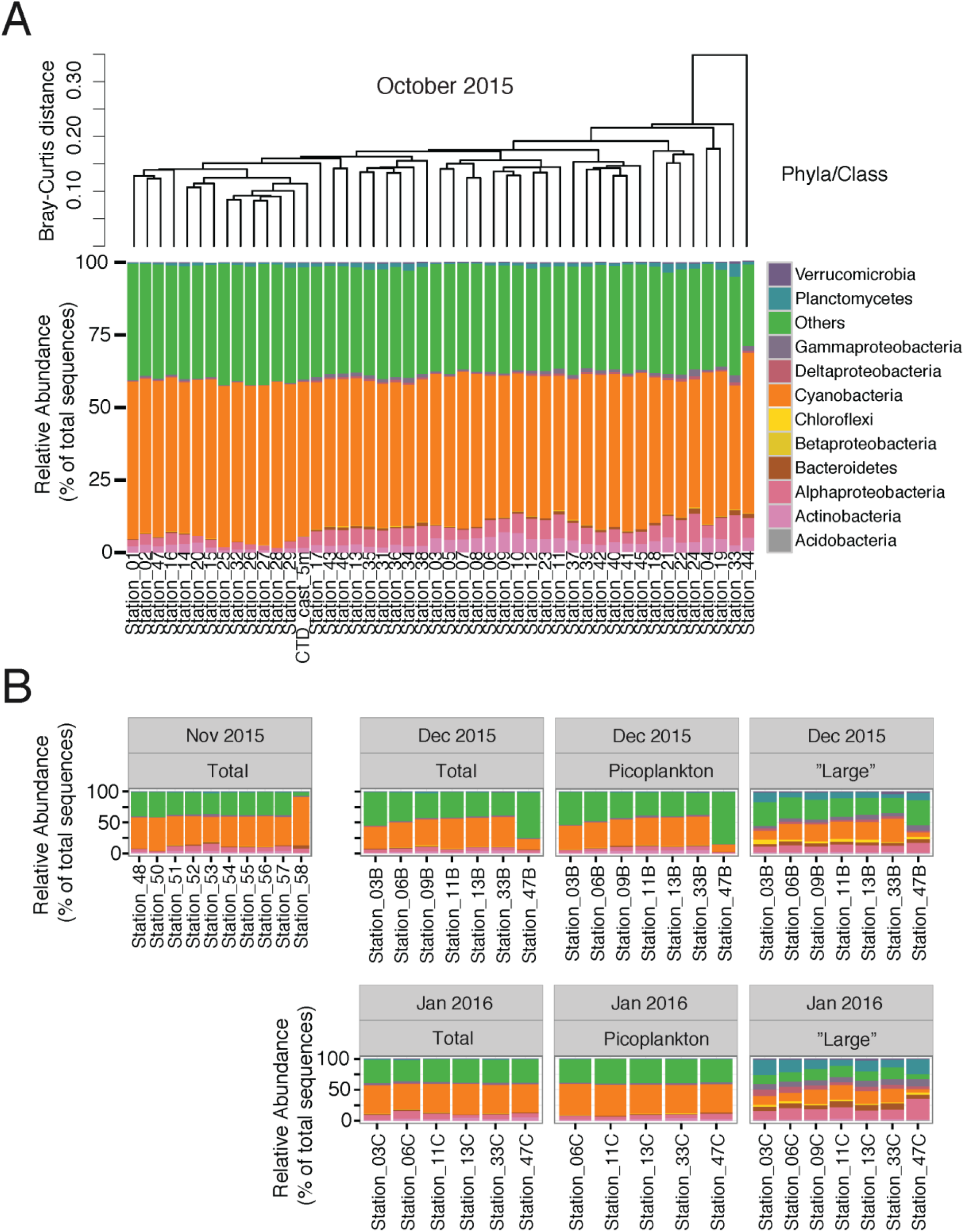
Variation in community composition during the October 2015 cruise (A), and November 2015, December 2015 and January 2016 cruises (B). Barcharts denote relative abundance (% of total sequences) among major bacterial groups at phyla/class. Dendrogram in (A) denote cluster analysis of variation in beta diversity estimated from Bray-Curtis distances. For the cruises in December 2015 and January 2016 in (B) total, picoplankton and “large” denote size-fractionated community DNA from >0.2 μm, >0.2 and <3.0 μm and 3.0 μm filters, respectively.

Among the top ten most abundant OTUs all were affiliated with *Prochlorococcus* sp., demonstrating homogenous relative abundances along the transect (Fig S2). Other less abundant OTUs that exhibited variance in relative abundance along the transect, included the OM-1 clade bacteria (*Candidatus* Actinomarina) and OTUs within the family *Rhodospirillaceae* affiliated with AEGEAN-169 marine group. Notably, the OM-1 clade bacteria appeared relatively enriched in a localized region north of Oahu (22° N, see arrows Fig. S3). Nevertheless, the variation in relative abundance among these OTUs was low.

## Occupancy-frequency distributions

The number of different species occupying different number of sites sampled during the transects were also examined to evaluate the shape of occupancy-frequency distributions. Bacterial communities sampled during the October 2015 cruise displayed a significant bimodal occupancy-frequency pattern. Most OTUs were found at a single site followed by a monotonical decrease in the number of OTUs occupying increasing number of sites but with a peak in the number of OTUs occupying all sites (Fig. 4A; Table S1). As this transect contained several samples obtained from around Station ALOHA this dataset were subsampled to only show stations 3, 6, 9, 13, 33 and 47 to keep a clear transect trajectory profile with distinct sites (see insert Fig. 4A). This subsampling exercise confirmed the significant bimodal pattern. In addition, samples from completely different stations in a trajectory 30 km Southeast of Honolulu were collected to validate the shape of occupancy-frequency distribution found in the Honolulu-Station ALOHA transect. Also for this transect a significant bimodal pattern were found (see insert Fig. 4A).

**Figure 4.**
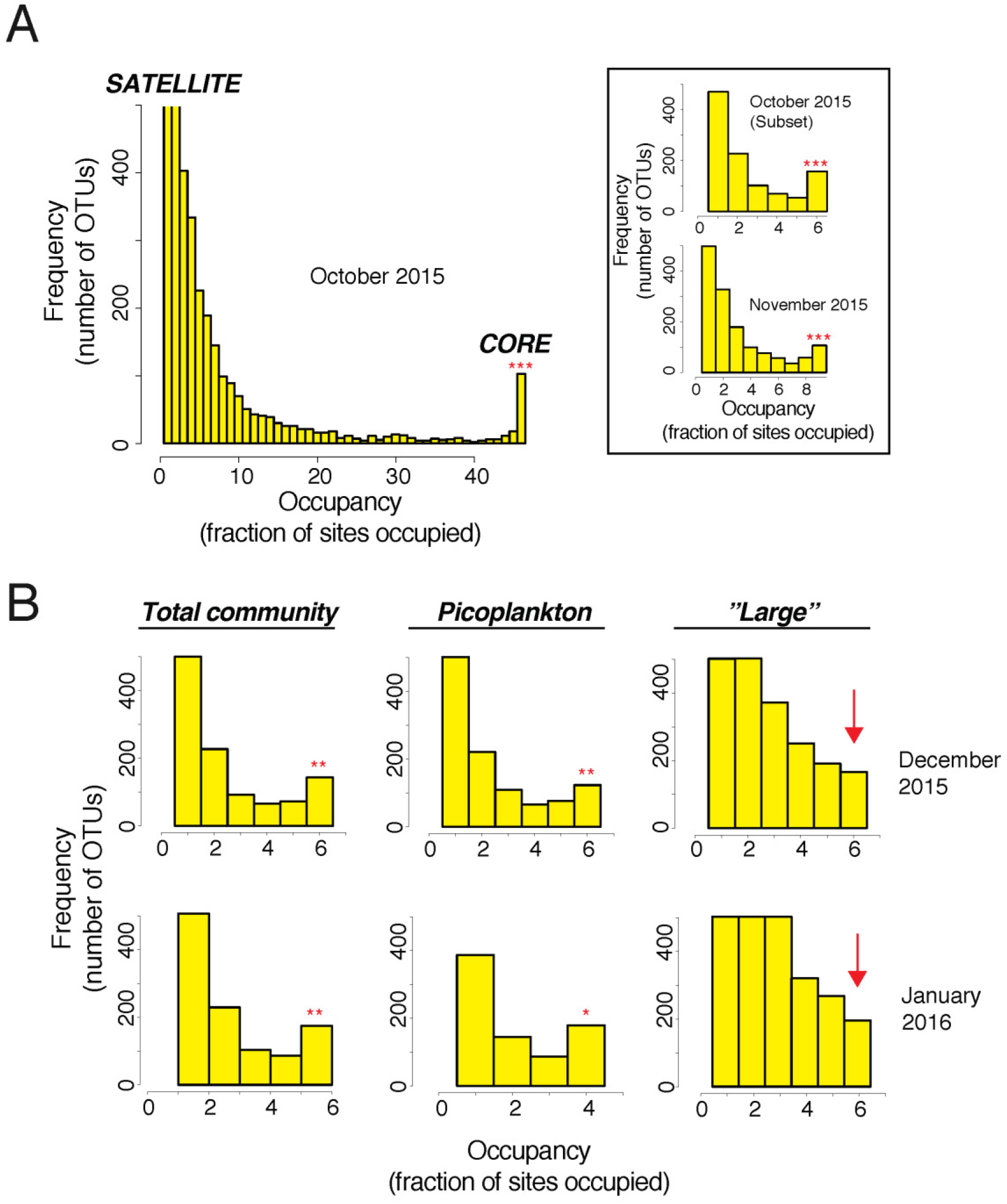
Core and satellite populations detected among occupancy-frequency distributions during the October 2015 cruise (A), and December 2015 and January 2016 cruises (B). Insert in (A) shows occupancy-frequency distributions of populations in a subset of the October 2015 cruise with the same stations sampled in the December 2015 and January 2016 cruises and the additional cruise in November 2015. Total, picoplankton and “large” denote size-fractionated community DNA from >0.2 μm, >0.2 and <3.0 μm and 3.0 μm filters, respectively. The y-axis is scaled at maximum 500 OTUs. Bimodality tests was performed using Mitchell-Olds and Shaw’s test for quadratic extremes (Mitchell-Olds and Shaw, 1987), a proxy for Tokeshi’s test (Tokeshi, 1992). ND=Not Determined. Significance level is indicated with “***”, “**” and “*” for p-values <0.001, <0.01 and <0.05, respectively.

The main hypotheses were that different parts of a bacterial community may exhibit different metapopulation dynamics; for example, taxa distributed among the picoplankton and larger bacterial assemblages might have different dispersal capabilities and be subjected to different environmental filters. To elucidate differences between such compartments the total community from the October 2015 cruise and size-fractionated communities from the December 2015 and January 2016 cruises were examined and taxa were analyzed at different taxonomic levels. Similar to the “total” community, the picoplankton assemblage demonstrated significant bimodal patterns (Fig. 4B; Table S1); in contrast, for the “large” bacterial size classes most OTUs were found at a single site followed by a monotonical decrease in the number of OTUs occupying increasing number of sites. Thus, in contrast to the picoplankton and total communities, the large size fractions exhibited unimodal occupancy-frequency patterns (Fig. 4B; Table S1). In addition, for particular OTUs binned at the phyla/class level detected in the October 2015 cruise varying occupancy-frequency distributions were observed. In fact, only Planctomycetes, Alphaproteobacteria, and Cyanobacteria contained OTUs that were found at all sites and only the latter two displayed bimodal patterns (Fig. S4; Table S1). Cyanobacteria had the most substantial bimodal pattern with a strong peak in number of OTUs occupying all sites. Moreover, most OTUs binned at the phyla/class level exhibited unimodal patterns for picoplankton and large size class communities (Fig. S5; Table S1).

More than 50% (64 out of 108) of the core OTUs detected from the October 2015 cruise was affiliated with *Prochlorococcus* sp. (Fig. 3; S6A). When analyzed separately, *Prochlorococcus* sp. OTUs exhibited a strong right-skewed occupancy-frequency distribution in the total and picoplankton size classes, with most OTUs occupying all sites. There were satellite phylotypes among the *Prochlorococcus* sp. affiliated OTUs detected at only a single site (Fig. S6). Notably, for the *Prochlorococcus* sp. OTUs in the large size class communities, neither unimodal nor bimodal occupancy-frequency patterns were detected (Fig. S6B). By analyzing phylogenetically distinct *Prochlorococcus* sp. OTUs and comparing those OTUs detected at all sites (core) vs. the OTUs only found in a single site (satellite) phylogenetic differences in core/satellite characteristics among closely related populations were examined. This analysis revealed that the most abundant core *Prochlorococcus* sp. OTUs were phylogenetically distinct from the most frequently observed satellite OTUs within the same genus (Fig. S7).

## Colonization and extinction rates

In the CSH (Hanski, 1982) predicted bimodal occupancy-frequency distributions of species based on stochastic variation in colonization and extinction rates following a quadratic relationship between these rates and occupancy. By calculating the number of OTUs successfully dispersed to new sites compared to the number of OTUs that disappeared from occupied sites colonization and extinction rates was estimated (Fig. 5). These calculations were performed using the December 2015 and January 2016 cruises where the same stations were sampled allowing for the observation of temporal changes in the number of occupied sites (d*P*/dt).

**Figure 5.**
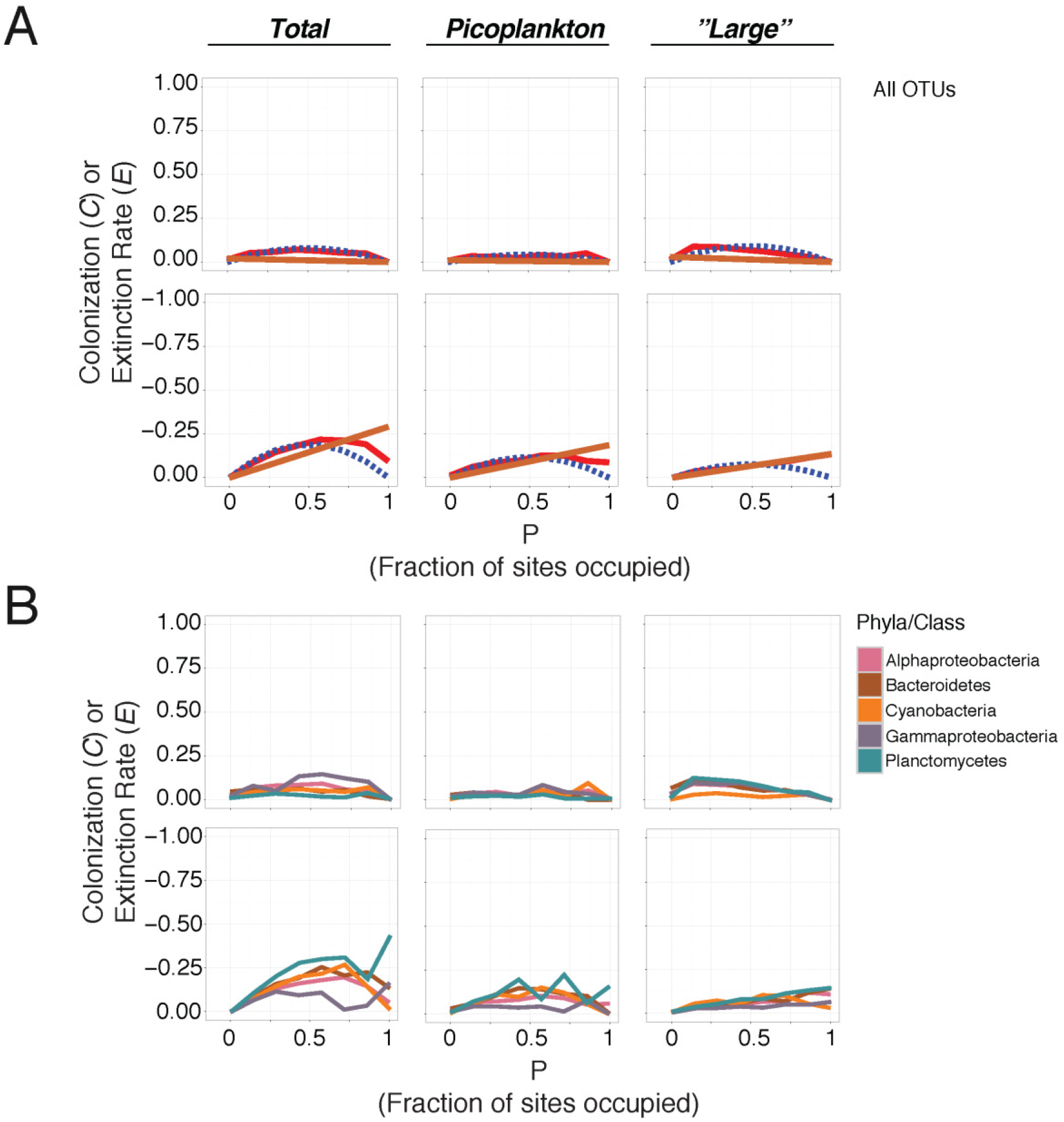
Measured colonization and extinction rates for all OTUs collectively (A), and for all OTUs within major bacterial groups and particular OTUs within the same genus (B). Observed rates in (A) were fitted with theoretical predictions from different metapopulation models.

The total bacterial community was characterized by a quadratic curve for both colonization and extinction rates. Notably, extinction rates were higher than colonization rates with a more distinct quadratic curve (Fig. 5A). Both colonization and extinction rates fitted well with the quadratic relationship described by the CSH (Hanski, 1982). For the picoplankton and large size class communities, the relationship between colonization rates and occupancy were more difficult to determine and none of the models tested resulted in robust fits to the observations (Fig. 5A). However, for the large plankton size class, extinction rates aligned well with the expectation of a linear relationship between extinction rates and occupancy as described in Levin’s model (Levin, 1974).

At the Phyla/Class level rates of colonization and extinction were plotted against occupancy for major bacterial lineages (Fig. 5B). Both Alphaproteobacteria and Cyanobacteria demonstrated quadratic relationships between extinction rates and occupancy in the total community. However, as above, colonization rates were more difficult to characterize. It was noteworthy that the extinction rates were both higher and more pronounced compared to colonization rates. Planctomycetes, Bacteroidetes and Gammaproteobacteria demonstrated curves that initially appeared to follow a quadratic relationship between extinction rates and occupancy, but either decreased followed by an increase in extinction rates with occupancy (Planctomycetes and Gammaproteobacteria) and/or did not decrease rapidly at maximum occupancy (Bacteroidetes). Unlike the patterns for all of the OTUs in the overall community analysis, resulting patterns among the picoplankton and large size class bacterial communities at phyla/class level were more difficult to validate using the different metapopulation models. Nevertheless, extinction rates among OTUs binned at phyla/class for the larger size class communities displayed a tendency toward a linear relationship between extinction rates and occupancy thus following Levin’s model (Levin, 1974) (Fig. 5B).

## Discussion

In the present paper metapopulation models were applied to describe bacterial community dynamics observed in the North Pacific Subtropical Gyre to elucidate mechanisms shaping biogeography. The results highlight that bimodal occupancy-frequency patterns are prominent in the North Pacific Subtropical Gyre but only among specific size classes and taxa of the bacterial communities. Total (≥0.2 μm size fraction) and picoplanktonic (≥0.2 μm ≤3.0 μm size fraction) communities typically displayed bimodal patterns, whereas the larger size class (≥3.0 μm filter fraction) community exhibited unimodal patterns. In instances where bimodal patterns were found quadratic relationships between rates of colonization/extinction and occupancy were observed. These findings indicate a strong positive feedback mechanism between local abundance and occupancy and the observed patterns generally fit Hanski’s metapopulation model, the CSH (Hanski, 1982). In agreement, significant bimodal occupancy-frequency distributions linked with quadratic colonization and extinction rates have been found among bacterial assemblages in the Baltic Sea Proper (Lindh et al., 2016).

## Environmental conditions and diversity regulate metapopulation dynamics

Concomitant measurements of environmental variables, bacterial abundance and ^3^H-leucine incorporation rates in the present paper allowed for linking mechanisms shaping biogeography with the prevailing environmental conditions and community functioning. Overall, the environment and community dynamics was highly stable throughout each of the transects performed coupled with core and satellite metapopulation dynamics. Bimodal patterns were typically coupled with lower community diversity and quadratic relationships between colonization/extinction rates and occupancy of OTUs in the total and picoplankton communities. In contrast, the large size fraction communities exhibited unimodal patterns linked with higher diversity and linear colonization and extinction rates of OTUs. These data suggest that positive feedbacks between local abundance and occupancy are important in structuring picoplankton bacterial communities when environmental conditions are homogenous and diversity is low. Still, it is noteworthy that the metadata in this study was derived from whole seawater and total community and the environmental conditions characterizing the size fractionated “large” communities are unknown. Further more focused studies coupling size fractionated community composition and functioning would be very rewarding for our understanding of metapopulation dynamics and ultimately provide a deeper mechanistic understanding of biogeography of both picoplankton and large size fraction communities.

## Metapopulation dynamics of Prochlorococcus sp

OTUs affiliated with *Prochlorococcus* sp. contributed up to 50 % of total sequences for the total and picoplankton communities. A majority of the detected core populations were in fact affiliated with *Prochlorococcus* sp., thus driving much of the observed metapopulation dynamics. In agreement with these findings, *Prochlorococcus* sp. is the dominant bacteria found in this system (Schmidt et al., 1991; Campbell and Vaulot, 1993; Eiler et al., 2011) and are ubiquitous in the oligotrophic ocean (Flombaum et al., 2013). However, this is the first study to examine metapopulation models for *Prochlorococcus* sp. and finding core- and satellite dynamics for this key organism. Although many of the *Prochlorococcus* sp. OTUs found in this study were distributed over all stations, OTUs within the same taxa had more restricted ranges geographically. Such OTUs with restricted distributions also added to the dominance of the left-skewed majority of rare satellite populations indicating dispersal and environmental filtering of particular *Prochlorococcus* ecotypes. Although this bacterium is ubiquitously distributed in the oligotrophic surface ocean, ecotypes within the same species have been suggested from variations in genome content and functional potential (Rocap et al., 2003). In fact, ecotypes of *Prochlorococcus* sp. have been observed in the Atlantic Ocean along environmental gradients with varying conditions implicating niche partitioning of closely related 16S rRNA sequences derived from the same species (Johnson et al., 2006). Single-cell genomic approaches have further emphasized that subpopulations of the same *Prochlorococcus* sp. are likely adapted to different environmental conditions (Kashtan et al., 2014). Taken together, *Prochlorococcus* sp. metapopulations exhibited core and satellite dynamics indicative of positive feedback mechanisms between local abundance and occupancy suggesting a rescue effect of core populations (Hanski, 1982; Hanski and Gyllenberg, 1993).

## Disentangling community compartments

Individual taxonomic groups and particular OTUs displayed different metapopulation dynamics suggesting that taxa likely have different dispersal capability and are subjected to environmental filtering. Further studies at deeper taxonomic levels are however warranted to examine dynamics of “large” communities at different spatial scales to determine the mechanisms that regulate their biogeographical distribution. From observing variation in the shape of occupancy-frequency distributions among complex assemblages studied in macroecology (Mehranvar and Jackson, 2001) and colleagues have suggested that pooling all taxa into the same metapopulation model obscured taxonomic differences. Such taxonomic differences in compartments of communities may ultimately be linked with different dispersal capabilities and assembly of distinct species (Lindström and Langenheder, 2012). Conclusions made on community structure from high-throughput sequencing of microbial assemblages coupled with community functioning should thus be done with care and relative to individual compartments.

## Acknowledgements

I acknowledge Matthew Church for valuable support and for providing comments on the paper, the HOT program science team for their assistance at sea, in the lab and for providing contextual data used in this study. In addition, I thank the captain, officers, and crew of the R/V *Ka‘imikai-O-Kanaloa* and *Kilo Moana* for their assistance on the cruises. I am also grateful to Lance Fujieki for his assistance with CTD data processing. Funding for this research derived from the Simons Collaboration on Ocean Processes and Ecology (SCOPE) to Matthew Church.

## Supporting information Supplementary Figures and Tables

**Figure S1.**
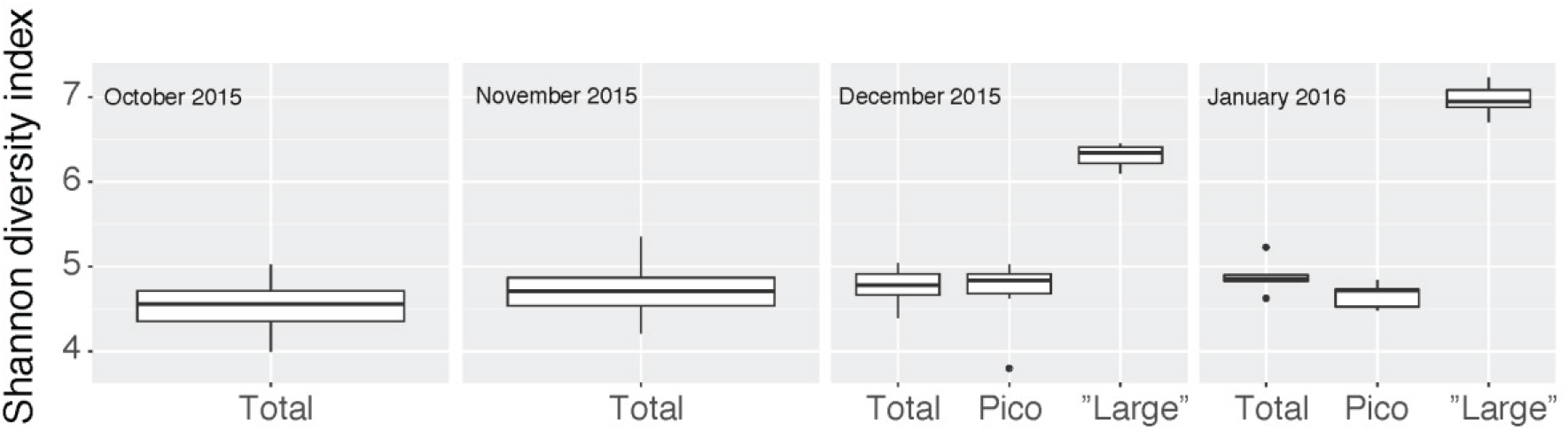
Variation in observed Shannon diversity index for all cruises.

**Figure S2.**
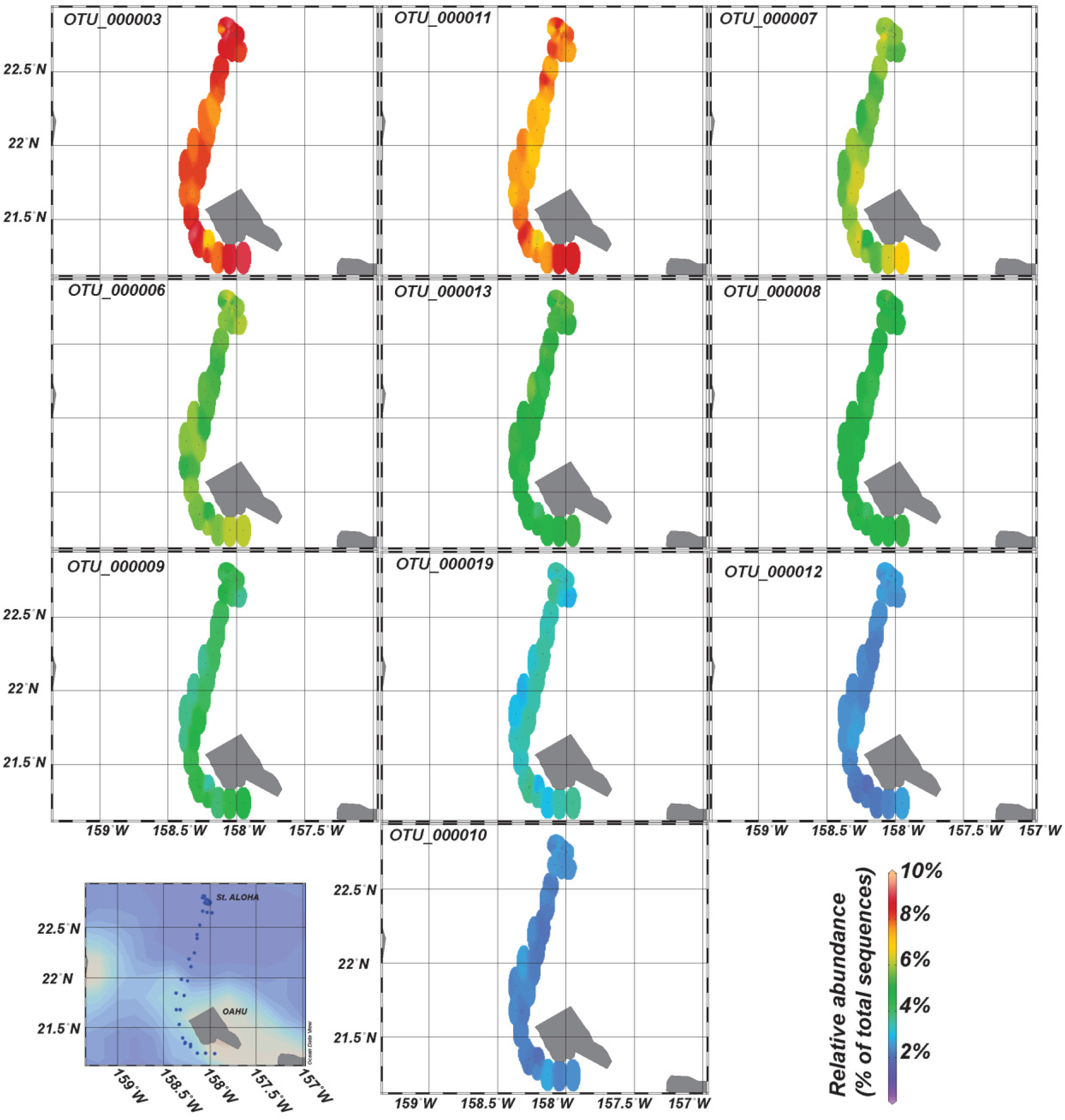
Relative abundances of the top ten most abundant OTUs found during the October 2015 transect. All OTUs were affiliated with *Prochlorococcus sp.* Colour shows interpolated relative abundances using the weighted-average gridding algorithm in Ocean Data View (http://odv.awi.de; version 4.7.8).

**Figure S3.**
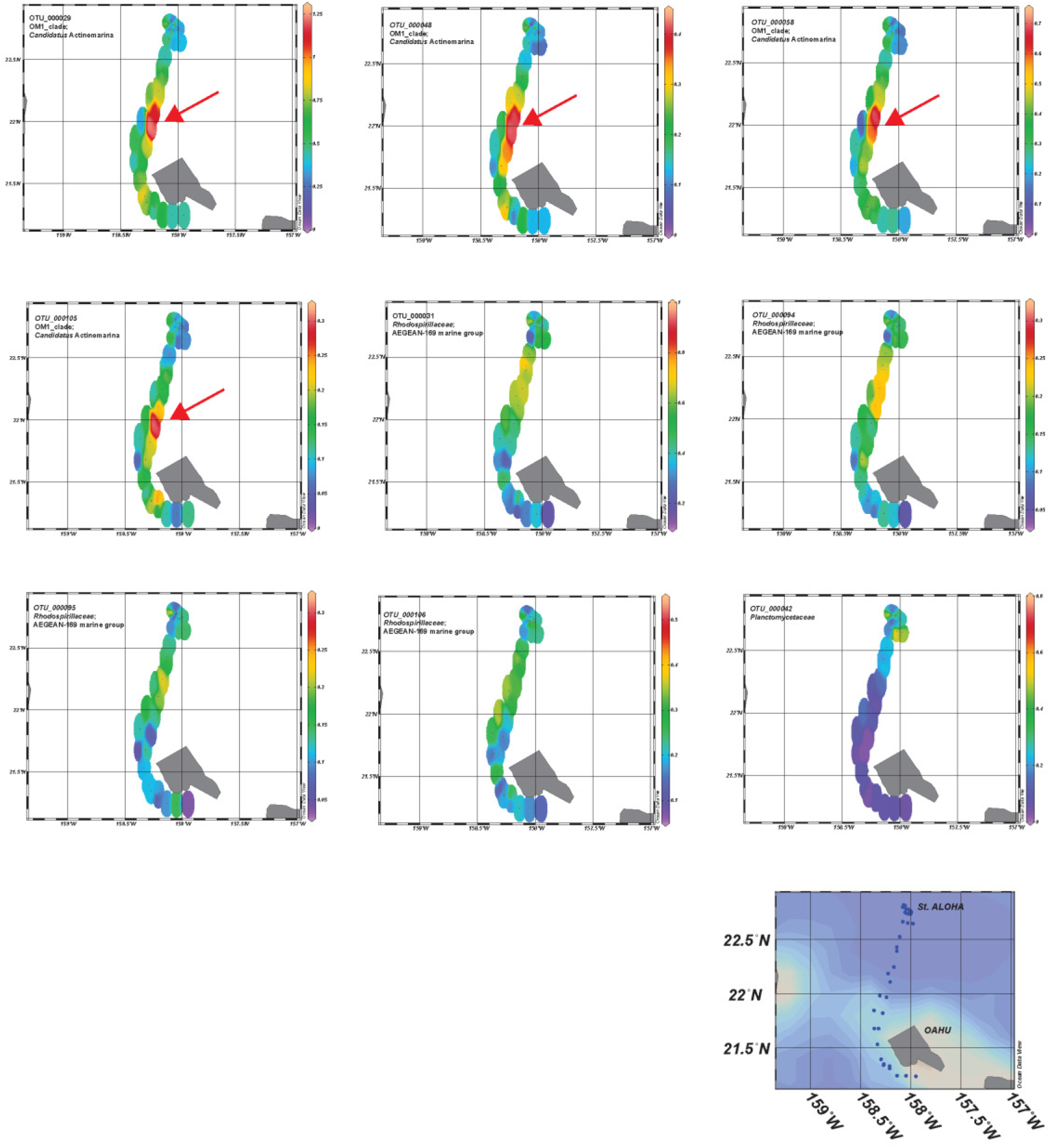
Relative abundances of the top ten OTUs exhibited the highest variance during the October 2015 transect. Colour shows interpolated relative abundances using the weighted-average gridding algorithm in Ocean Data View (http://odv.awi.de; version 4.7.8). Arrows denote hotspots in relative abundance detected for OM-1 clade *Candidatus* Actinomarina.

**Figure S4.**
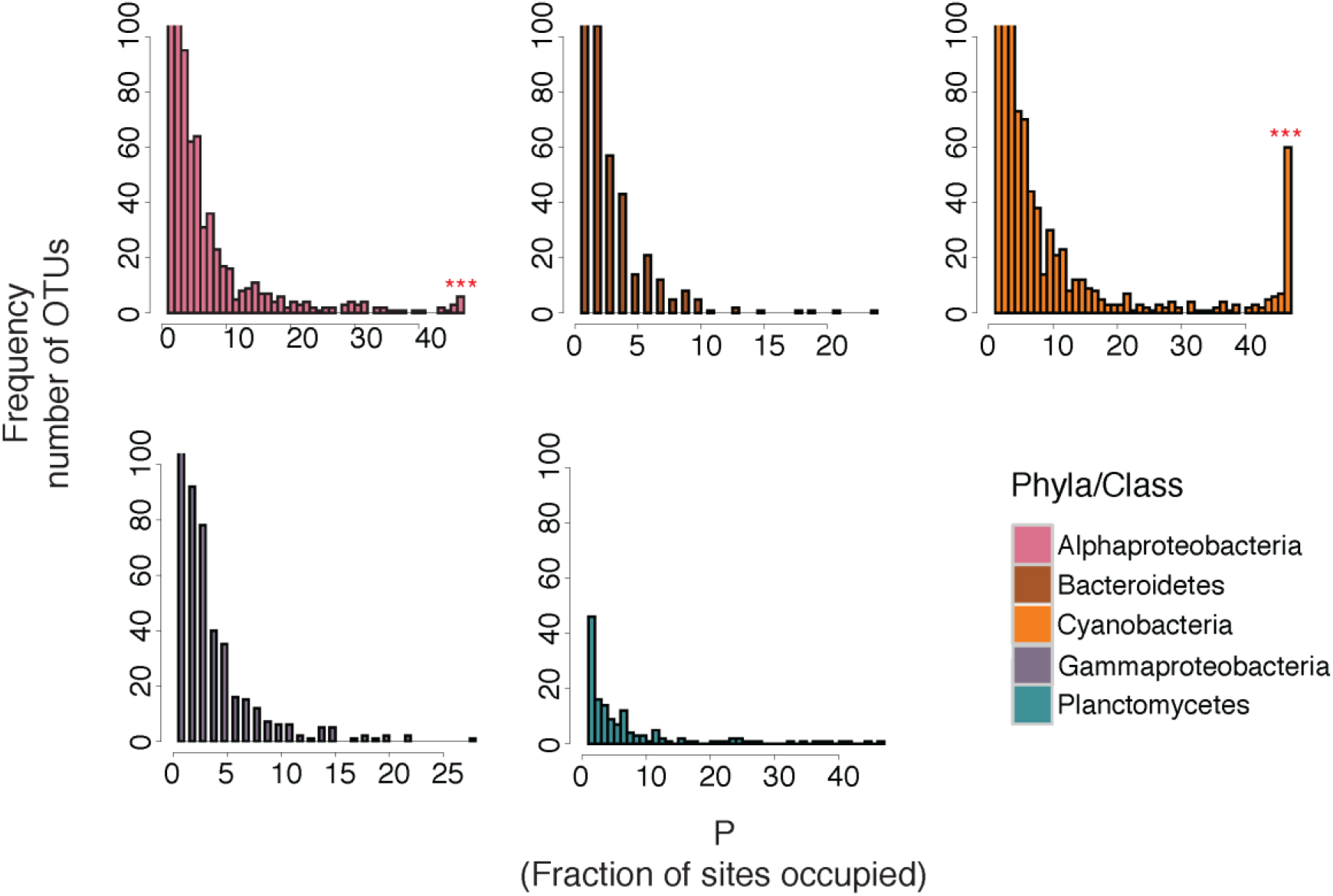
Occupancy-frequency distributions of populations binned by phyla/class during the October 2015 cruise. Bimodality tests was performed using Mitchell-Olds and Shaw’s test for quadratic extremes (Mitchell-Olds and Shaw, 1987), a proxy for Tokeshi’s test (Tokeshi, 1992). ND=Not Determined. Significance level is indicated with “***”, “**” and “*” for p-values <0.001, <0.01 and <0.05, respectively.

**Figure S5.**
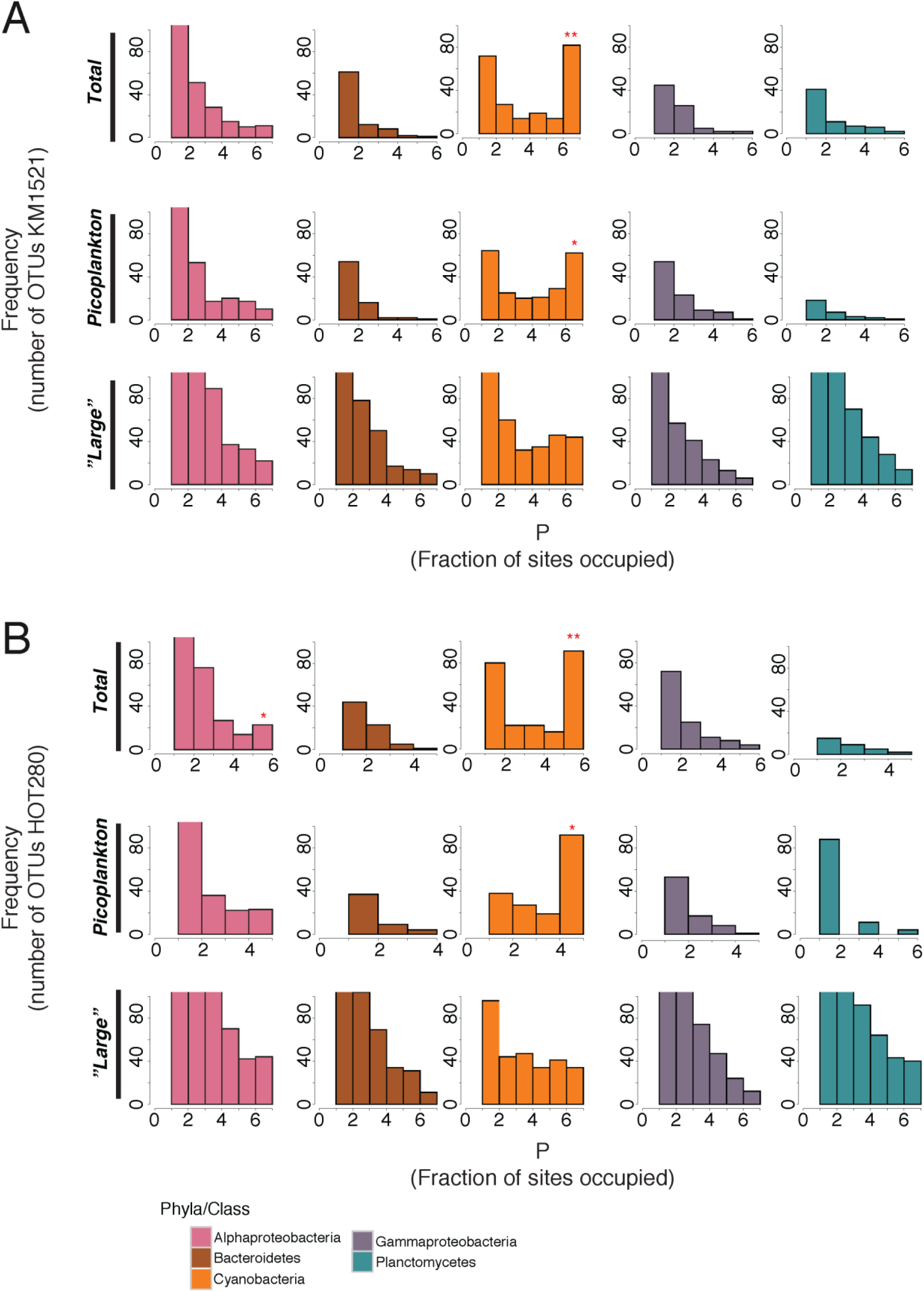
Occupancy-frequency distributions of populations binned by phyla/class during the December 2015 and January 2016 cruises. Total, picoplankton and “large” denote size-fractionated community DNA from >0.2 μm, >0.2 and <3.0 μm and 3.0 μm filters, respectively. Bimodality tests was performed using Mitchell-Olds and Shaw’s test for quadratic extremes (Mitchell-Olds and Shaw, 1987), a proxy for Tokeshi’s test (Tokeshi, 1992). ND=Not Determined. Significance level is indicated with “***”, “**” and “*” for p-values <0.001, <0.01 and <0.05, respectively.

**Figure S6.**
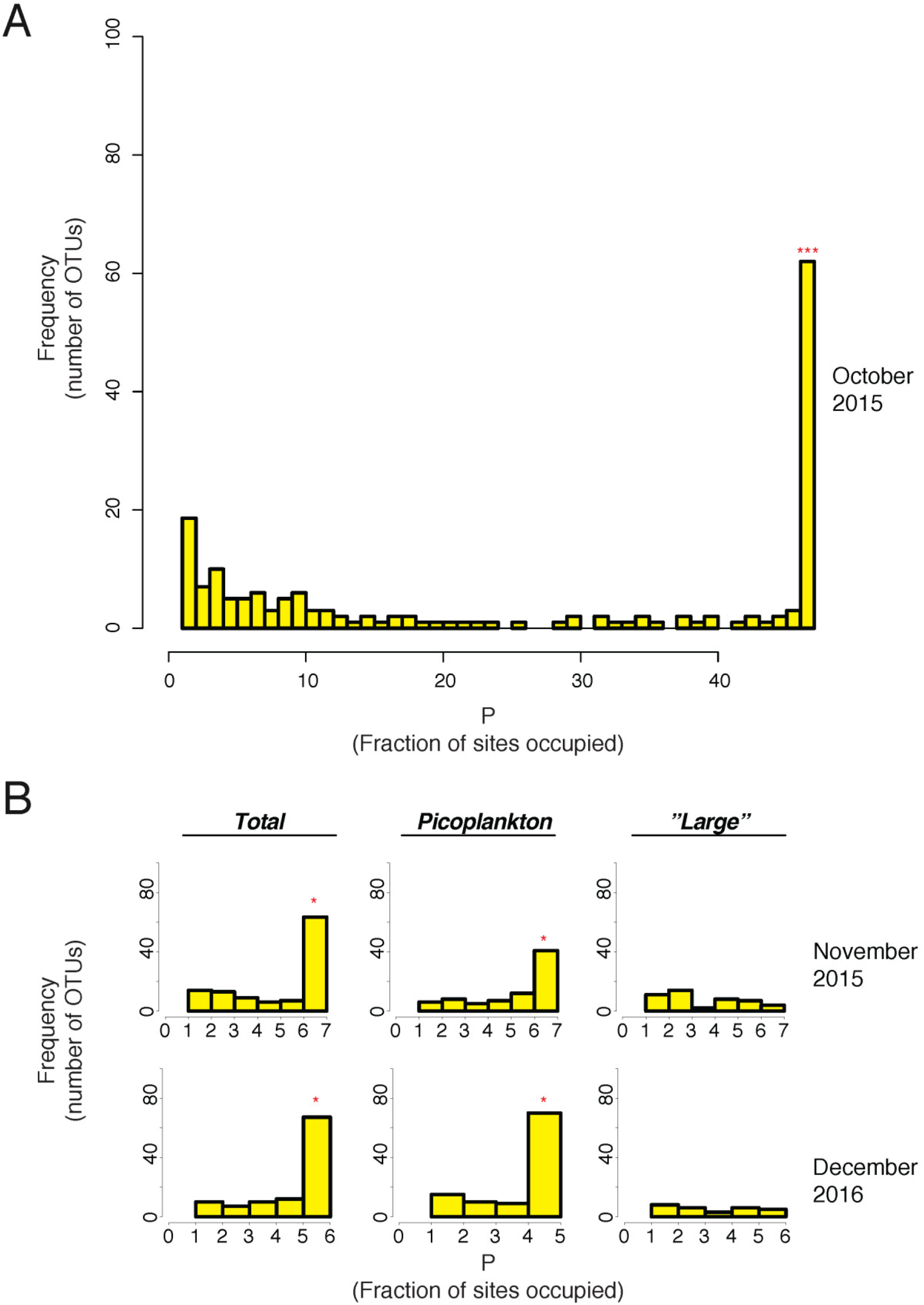
Occupancy-frequency distributions of OTUs affiliated with *Prochlorococcus* sp. during the October 2015 (A), and December 2015 and January 2016 cruises (B). Total, picoplankton and “large” in (B) denote size-fractionated community DNA from >0.2 μm, >0.2 and <3.0 μm and >3.0 μm filters, respectively. Bimodality tests was performed using Mitchell-Olds and Shaw’s test for quadratic extremes (Mitchell-Olds and Shaw, 1987), a proxy for Tokeshi’s test (Tokeshi, 1992). ND=Not Determined. Significance level is indicated with “***”, “**” and”*” for p-values <0.001, <0.01 and <0.05, respectively.

**Figure S7.**
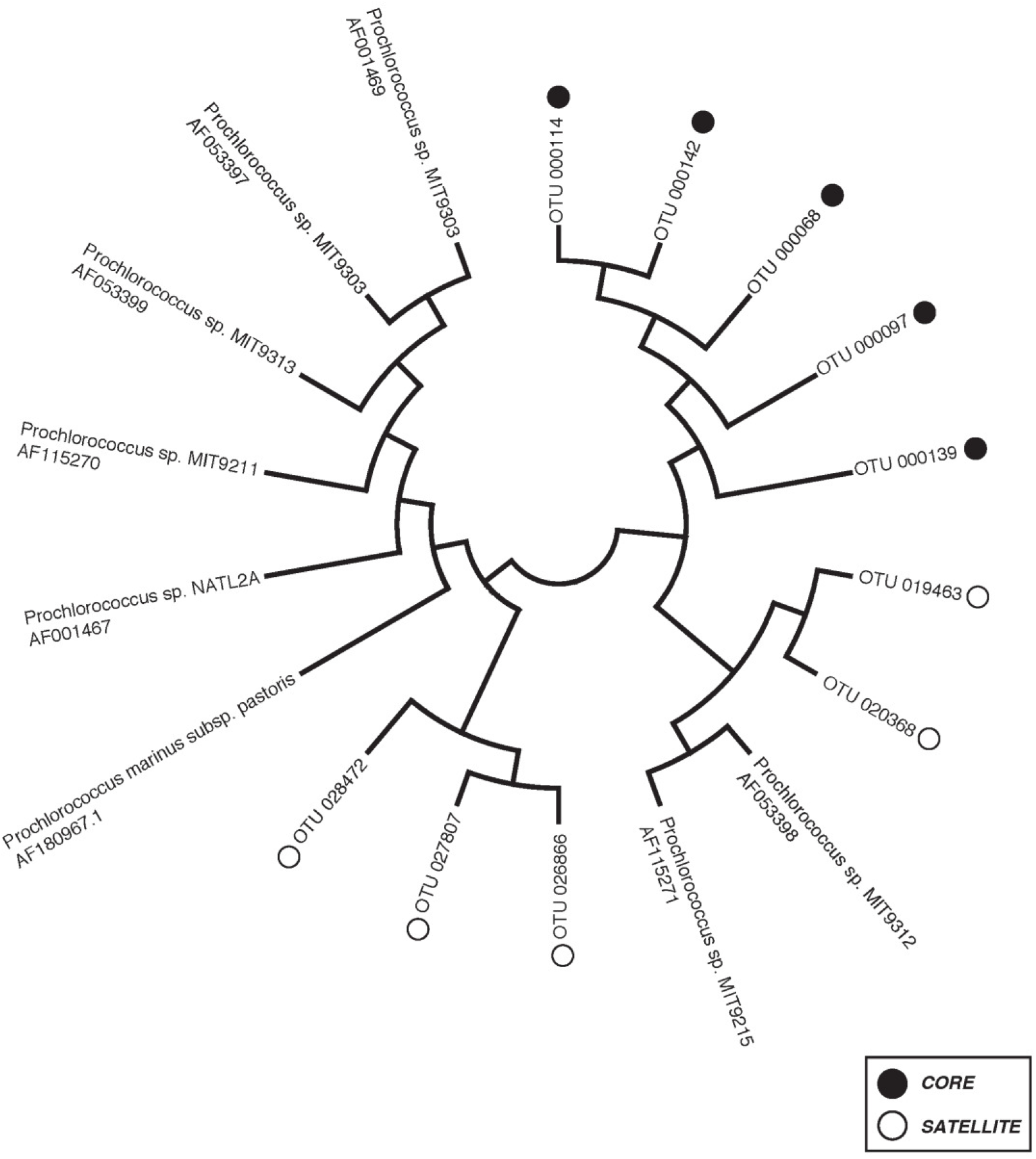
Maximum-likelihood tree of OTUs affiliated with *Prochlorococcus* sp. exhibiting core or satellite metapopulation dynamics obtained from 16S rRNA gene data from all cruises. The evolutionary history was inferred by using the Maximum Likelihood method based on the Tamura-Nei model (Tamura and Nei, 1993). The bootstrap consensus tree inferred from 999 replicates is taken to represent the evolutionary history of the taxa analyzed (Felsenstein, 1985). Branches corresponding to partitions reproduced in less than 50% bootstrap replicates are collapsed. Initial tree(s) for the heuristic search were obtained automatically by applying Neighbor-Join and BioNJ algorithms to a matrix of pairwise distances estimated using the Maximum Composite Likelihood (MCL) approach, and then selecting the topology with superior log likelihood value. The analysis involved 18 nucleotide sequences. All positions containing gaps and missing data were eliminated. There were a total of 427 positions in the final dataset. Evolutionary analyses were conducted in MEGA7 (Kumar et al., 2016).

**Table S1.**
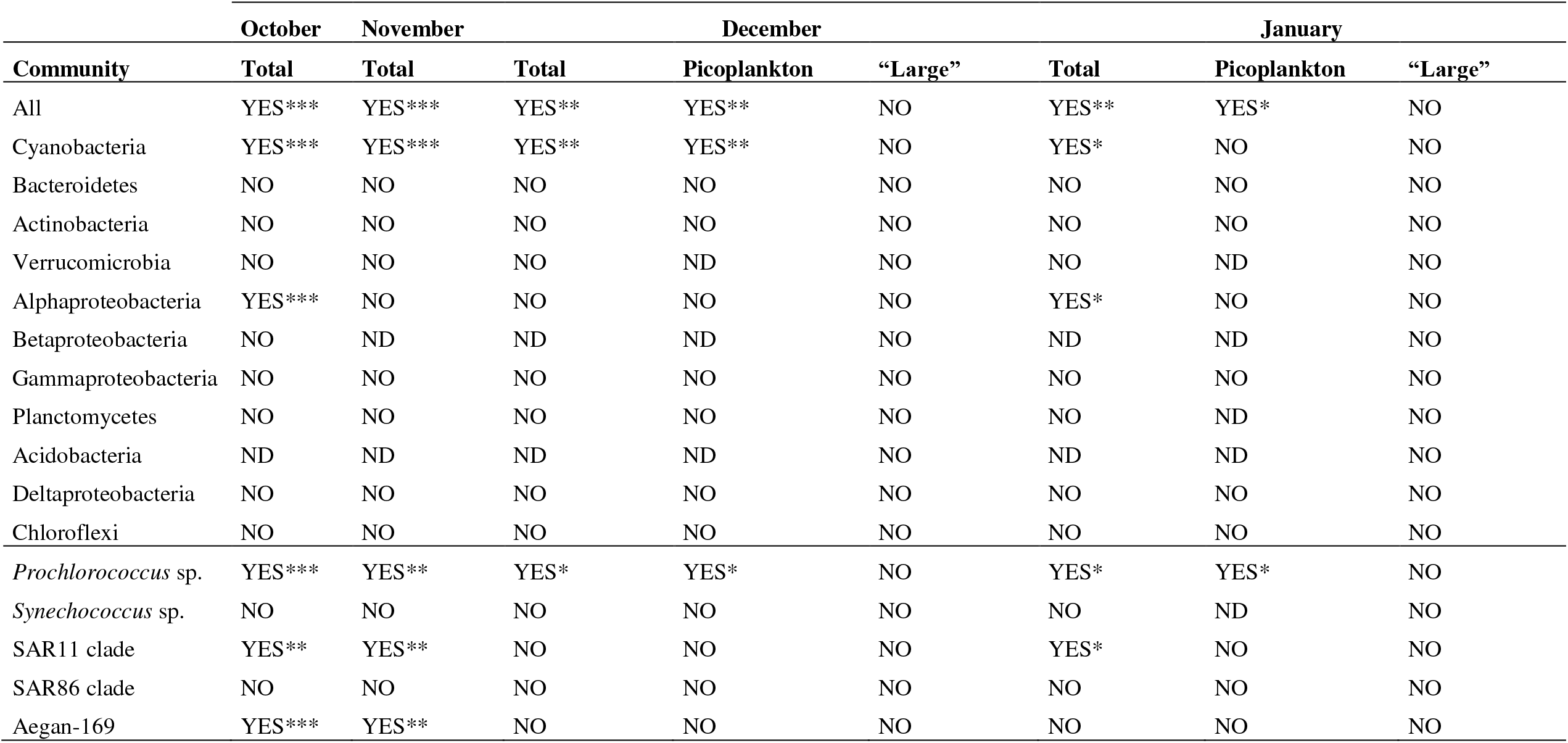
Prevalence of significantly bimodal occupancy-frequency patterns in different community compartments (size fractions and taxa) in the present paper. Bimodality tests was performed using Mitchell-Olds and Shaw’s test for quadratic extremes (Mitchell-Olds and Shaw, 1987), a proxy for Tokeshi’s test (Tokeshi, 1992). ND=Not Determined. Significance level is indicated with “***”, “**” and “*” for p-values <0.001, <0.01 and <0.05, respectively.

## References

Alonso-Saez, L., Diaz-Perez, L., and Moran, X.A. (2015) The hidden seasonality of the rare biosphere in coastal marine bacterioplankton. Environ Microbiol 17: 3766–3780.

Brown, J.H. (1984) On the relationship between abundance and distribution of species American Naturalist 124: 255–279.

Campbell, L., and Vaulot, D. (1993) Photosynthetic picoplankton community structure in the subtropical North Pacific Ocean near Hawaii (station ALOHA). Deep Sea Res Pt I 40: 2043–2060.

Crump, B.C., Hopkinson, C.S., Sogin, M.L., and Hobbie, J.E. (2004) Microbial Biogeography along an Estuarine Salinity Gradient: Combined Influences of Bacterial Growth and Residence Time. Appl Environ Microbiol 70: 1494–1505.

Edgar, R.C. (2013) UPARSE: highly accurate OTU sequences from microbial amplicon reads. Nat Methods 10: 996–998.

Eiler, A., Hayakawa, D., and Rappé, M. (2011) Non-Random Assembly of Bacterioplankton Communities in the Subtropical North Pacific Ocean. Front Microbiol 2.

Felsenstein, J. (1985) Confidence Limits on Phylogenies: An Approach Using the Bootstrap. Evolution 39: 783–791.

Flombaum, P., Gallegos, J.L., Gordillo, R.A., Rincón, J., Zabala, L.L., Jiao, N. et al. (2013) Present and future global distributions of the marine Cyanobacteria Prochlorococcus and Synechococcus. Proc Natl Acad Sci USA 110: 9824–9829.

Fuhrman, J.A., Hewson, I., Schwalbach, M.S., Steele, J.A., Brown, M.V., and Naeem, S. (2006) Annually reoccurring bacterial communities are predictable from ocean conditions. Proc Natl Acad Sci USA 103: 13104–13109.

Galand, P.E., Potvin, M., Casamayor, E.O., and Lovejoy, C. (2010) Hydrography shapes bacterial biogeography of the deep Arctic Ocean. ISME J 4: 564–576.

Ghiglione, J.F., Galand, P.E., Pommier, T., Pedros-Alio, C., Maas, E.W., Bakker, K. et al. (2012) Pole-to-pole biogeography of surface and deep marine bacterial communities. Proc Natl Acad Sci USA 109: 17633–17638.

Gotelli, N.J. (1991) Metapopulation models: the rescue effect, the propagule rain, and the core-satellite hypothesis. Am Nat X: 768–776.

Hanski, I. (1982) Dynamics of regional distribution - the core and satellite species hypothesis. Oikos 38: 210–221.

Hanski, I., and Gyllenberg, M. (1993) Two General Metapopulation Models and the Core-Satellite Species Hypothesis. Am Nat 142: 17–41.

Hanson, C.A., Fuhrman, J.A., Horner-Devine, M.C., and Martiny, J.B. (2012) Beyond biogeographic patterns: processes shaping the microbial landscape. Nat Rev Microbiol 10: 497–506.

Hercos, A.P., Sobansky, M., Queiroz, H.L., and Magurran, A.E. (2013) Local and regional rarity in a diverse tropical fish assemblage. Paper Vol: Page.

Herlemann, D.P., Labrenz, M., Jurgens, K., Bertilsson, S., Waniek, J.J., and Andersson, A.F. (2011) Transitions in bacterial communities along the 2000 km salinity gradient of the Baltic Sea. ISME J Vol: Page.

Hubbell, S.P. (2001) The unified neutral theory of biodiversity and biogeography (MPB-32): Princeton University Press.

Hugerth, L.W., Wefer, H.A., Lundin, S., Jakobsson, H.E., Lindberg, M., Rodin, S. et al. (2014) DegePrime, a program for degenerate primer design for broad-taxonomic-range PCR in microbial ecology studies. Appl Environ Microbiol 80: 5116–5123.

Johnson, Z.I., Zinser, E.R., Coe, A., McNulty, N.P., Woodward, E.M.S., and Chisholm, S.W. (2006) Niche Partitioning Among Prochlorococcus Ecotypes Along Ocean-Scale Environmental Gradients. Science 311: 1737–1740.

Kashtan, N., Roggensack, S.E., Rodrigue, S., Thompson, J.W., Biller, S.J., Coe, A. et al. (2014) Single-Cell Genomics Reveals Hundreds of Coexisting Subpopulations in Wild Prochlorococcus. Science 344: 416–420.

Kirchman, D., K′Ness, E., and Hodson, R. (1985) Leucine incorporation and its potential as a measure of protein synthesis by bacteria in natural aquatic systems. Appl Environ Microbiol 49: 599–607.

Kirchman, D.L., Dittel, A.I., Malmstrom, R.R., and Cottrell, M.T. (2005) Biogeography of major bacterial groups in the Delaware Estuary. Limn Oceanogr 50: 1697–1706.

Kumar, S., Stecher, G., and Tamura, K. (2016) MEGA7: Molecular Evolutionary Genetics Analysis version 7.0 for bigger datasets. Molecular Biology and Evolution.

Leibold, M.A., Holyoak, M., Mouquet, N., Amarasekare, P., Chase, J.M., Hoopes, M.F. et al. (2004) The metacommunity concept: a framework for multi-scale community ecology. Ecol Lett 7: 601–613.

Levin, S.A. (1974) Dispersion and population interactions Am Nat 108: 207–228.

Lindh, M.V., Figueroa, D., Sjöstedt, J., Baltar, F., Lundin, D., Andersson, A. et al. (2015) Transplant experiments uncover Baltic Sea basin-specific responses in bacterioplankton community composition and metabolic activities. Front Microbiol 6: Page.

Lindh, M.V., Sjöstedt, J., Ekstam, B., Casini, M., Lundin, D., Hugerth, L.W. et al. (2016) Metapopulation theory identifies biogeographical patterns among core and satellite marine bacteria scaling from tens to thousands of kilometers. Environ Microbiol Vol: Page.

Lindström, E.S., and Langenheder, S. (2012) Local and regional factors influencing bacterial community assembly. Environ Microbiol Rep 4: 1–9.

Lindström, E.S., Feng, X.M., Graneli, W., and Kritzberg, E.S. (2010) The interplay between bacterial community composition and the environment determining function of inland water bacteria. Limnol Oceanogr 55: 2052–2060.

Martiny, J.B.H., Bohannan, B.J.M., Brown, J.H., Colwell, R.K., Fuhrman, J.A., Green, J.L. et al. (2006) Microbial biogeography: putting microorganisms on the map. Nat Rev Microbiol 4: 102–112.

McGeoch, M.A., and Gaston, K.J. (2002) Occupancy frequency distributions: patterns, artefacts and mechanisms. Biol Rev 77: 311–331.

Mehranvar, L., and Jackson, D.A. (2001) History and taxonomy: their roles in the coresatellite hypothesis. Oecologia 127: 131–142.

Mitchell-Olds, T., and Shaw, R.G. (1987) Regression Analysis of Natural Selection: Statistical Inference and Biological Interpretation. Evolution 41: 1149–1161.

Mouquet, N., and Loreau M (2002) Coexistence in Metacommunities: The Regional Similarity Hypothesis. Am Nat 159: 420–426.

Oksanen, J., Guillaume Blanchet, F., Kindt, R., Legendre, Pierre., O′Hara, R. B., Simpson G. L., et al. (2010) vegan: Community Ecology Package. R package version 1.17–5. https://cran.r-project.org/web/packages/vegan/index.html.

Pace, M.L., del Giorgio, P., Fischer, D., Condon, R., and Malcom, H. (2004) Estimates of bacterial production using the leucine incorporation method are influenced by differences in protein retention of microcentrifuge tubes. Limnol Oceanogr Meth 2: 55–61.

Poisot, T., Pequin, B., and Gravel, D. (2013) High-throughput sequencing: a roadmap toward community ecology. Ecol Evol 3: 1125–1139.

Pommier, T., Canback, B., Riemann, L., Bostrom, K.H., Simu, K., Lundberg, P. et al. (2007) Global patterns of diversity and community structure in marine bacterioplankton. Mol Ecol 16: 867–880.

Purdy, K.J., Hurd, P.J., Moya-Larano, J., Trimmer, M., Oakley, B.B., Woodward, G., and Guy, W. (2010) Systems biology for ecology: from molecules to ecosystems. Adv Ecol Res 43: 87–149.

Quast, C., Pruesse, E., Yilmaz, P., Gerken, J., Schweer, T., Yarza, P. et al. (2013) The SILVA ribosomal RNA gene database project: improved data processing and web-based tools. Nucl Acids Res 41: D590–596.

R Core Team. (2014) R: A Language and Environment for Statistical Computing. https://cran.r-project.org/

Rocap, G., Larimer, F.W., Lamerdin, J., Malfatti, S., Chain, P., Ahlgren, N.A. et al. (2003) Genome divergence in two Prochlorococcus ecotypes reflects oceanic niche differentiation. Nature 424: 1042–1047.

Salazar, G., Cornejo-Castillo, F.M., Benitez-Barrios, V., Fraile-Nuez, E., Alvarez-Salgado, X.A., Duarte, C.M. et al. (2016) Global diversity and biogeography of deep-sea pelagic prokaryotes. ISME J 10: 596–608.

Schmidt, T.M., DeLong, E.F., and Pace, N.R. (1991) Analysis of a marine picoplankton community by 16S rRNA gene cloning and sequencing. J Bacteriol 173: 4371–4378.

Smith, D.C., and Azam, F. (1992) A simple, economical method for measuring bacterial protein synthesis rates in seawater using 3H-leucine. Marine Microb Food Webs 6: 107–114.

Sunagawa, S., Coelho, L.P., Chaffron, S., Kultima, J.R., Labadie, K., Salazar, G. et al. (2015) Structure and function of the global ocean microbiome. Science 348: Page.

Tamura, K., and Nei, M. (1993) Estimation of the number of nucleotide substitutions in the control region of mitochondrial DNA in humans and chimpanzees. Mol Biol Evol 10: 512–526.

Tokeshi, M. (1992) Dynamics of distribution in animal communities - theory and analysis. Res Pop Ecol 34: 249–273.

Unterseher, M., Jumpponen, A.R.I., ÖPik, M., Tedersoo, L., Moora, M., Dormann, C.F., and Schnittler, M. (2011) Species abundance distributions and richness estimations in fungal metagenomics – lessons learned from community ecology. Mol Ecol 20: 275–285.

van Rensburg, B.J., McGeoch, M.A., Matthews, W., Chown, S.L., and van Jaarsveld, A.S. (2000) Testing generalities in the shape of patch occupancy frequency distributions Ecology 81: 3163–3177.

Verberk, W.C.E.P., Van Der Velde, G., and Esselink, H. (2010) Explaining abundance – occupancy relationships in specialists and generalists: a case study on aquatic macroinvertebrates in standing waters. J Anim Ecol 79: 589–601.

Viviani, D.A., and Church, M.J. (2017) Decoupling between bacterial production and primary production over multiple time scales in the North Pacific Subtropical Gyre. Deep Sea Res Pt I 121: 132–142.

Wardle, D.A., Bardgett, R.D., Callaway, R.M., and Van der Putten, W.H. (2011) Terrestrial Ecosystem Responses to Species Gains and Losses. Science 332: 1273–1277.

Wickham, H. (2009) ggplot2: elegant graphics for data analysis. New York: Springer.

Östman, O., Drakare, S., Kritzberg, E.S., Langenheder, S., Logue, J.B., and Lindström, E.S. (2010) Regional invariance among microbial communities. Ecol Lett 13: 118–127.

